# IFN-λ is protective against lethal oral *Toxoplasma gondii* infection

**DOI:** 10.1101/2023.02.24.529861

**Authors:** Mateo Murillo-León, Aura M. Bastidas-Quintero, Niklas S. Endres, Daniel Schnepf, Estefanía Delgado-Betancourt, Annette Ohnemus, Gregory A. Taylor, Martin Schwemmle, Peter Staeheli, Tobias Steinfeldt

## Abstract

Interferons are essential for innate and adaptive immune responses against a wide variety of pathogens. Interferon lambda (IFN-λ) protects mucosal barriers during pathogen exposure. The intestinal epithelium is the first contact site for *Toxoplasma gondii* (*T. gondii*) with its hosts and the first defense line that limits parasite infection. Knowledge of very early *T. gondii* infection events in the gut tissue is limited and a possible contribution of IFN-λ has not been investigated so far. Here, we demonstrate with systemic interferon lambda receptor (IFNLR1) and conditional (Villin-Cre) knockout mouse models and bone marrow chimeras of oral *T. gondii* infection and mouse intestinal organoids a significant impact of IFN-λ signaling in intestinal epithelial cells and neutrophils to *T. gondii* control in the gastrointestinal tract. Our results expand the repertoire of interferons that contribute to the control of *T. gondii* and may lead to novel therapeutic approaches against this world-wide zoonotic pathogen.

## Introduction

Interferons (IFNs) are essential regulators of the host immune response against a variety of microbial infections. Depending on the nature of the unique cell surface receptors required for signal transduction, they are classified into type I interferons (17 in humans and 18 in mice) that bind to the heterodimeric receptor complex consisting of IFN-α/β receptor 1 (IFNAR1) and IFN-α/β receptor 2 (IFNAR2) heterodimers^3^, type II interferon (IFN-γ) that binds to IFN-γ receptor 1 (IFNGR1) and IFN-γ receptor 2 (IFNGR2) heterodimers^4^, and type III interferons (IFN-λ1-4 in humans and IFN-λ2/3 in mice^5^) that bind to IFN-λ receptor 1 (IFNLR1) and IL10 receptor subunit β (IL10RB) heterodimers^6–8^.

Due to the high degree of overlapping downstream signaling^7,8^, the IFN type I and type III systems were initially considered to be functionally redundant^6,7^. However, in recent years, unique features of the IFN-λ-mediated immune response against respiratory and gastrointestinal viruses^2,9–13^, fungi^14^, bacteria^15,16^ and parasites^17,18^ have been demonstrated. While almost all nucleated cells respond to type I and type II interferons, the function of IFN-λ is primarily restricted to epithelial cells at barrier surfaces^2,9–13,19,20^ and some immune cell types^14,21–25^ due to the tissue tropism of IFNLR1^13^. As an exception, intestinal epithelial cells of adult mice do not express a functional type I IFN receptor and therefore strongly rely on the IFN-λ system for antimicrobial defense^11,26^ Depending on the cell type, the downstream signaling of the type I and III IFN system can differ significantly. Especially in neutrophils, a subset of inflammatory cytokines is induced by type I but not by type III signaling^21,23,27^. IFN-λ is therefore believed to act locally as the first line of defense against invading pathogens on mucosal surfaces possibly without activating the detrimental immune responses mediated by IFN type I^28,1,13^. IFN-λ has also been described to enhance adaptive immune responses at these sites^2,29^.

*Toxoplasma gondii* (*T. gondii*) is a foodborne obligate intracellular parasite related to the *Plasmodium* genus. About 25-30 % of the world human population is infected but local seroprevalences can vary significantly^30^. Mild “flu-like” symptoms may occur upon infection for several weeks or months. In patients with a compromised immune system on the other hand, the parasite can cause serious health problems. Transmission to the fetus upon primary infection of the mother may lead to miscarriage, stillbirth or child disability^31^. The natural route of infection with *T. gondii* is the uptake of infective stages, either contained in tissue cysts (bradyzoites) of intermediate hosts or oocysts (sporozoites) released into the environment by all members within the family of *Felidae*^32,33^. After ingestion, once tissue cysts or oocysts reach the small intestine, released parasites can cross the intestinal epithelial barrier (IEB) by either paracellular transmigration or penetration of the apical cell membrane and passing through the basolateral side to reach the underlying lamina propria^33,34^ In addition, neutrophils that transmigrate to the intestinal lumen after oral *T. gondii* infection, are hijacked by the parasite in order to be spread across the intestine and are found preferentially infected by *T. gondii* in the lamina propria^35^.

Because of the sympatry of cats and mice, a mouse model of toxoplasmosis is of medical importance for human infections. The innate and adaptative immune responses against *T. gondii* rely on IFN-γ that is produced early after infection by natural killer (NK) cells, neutrophils and T cells^36–39^. Two families of IFN-γ-inducible GTPases are paramount for innate immunity against *T. gondii* in mice, the Immunity-Related GTPases (IRG proteins) and Guanylate Binding Proteins (GBP proteins)^40,41^. Certain family members were demonstrated to accumulate at the parasitophorous vacuole membrane (PVM) of *T. gondii*, a prerequisite for subsequent membrane disintegration and parasite death^42^. Type I IFNs have also been shown to play a protective role during *T. gondii* infection by limiting the growth of parasite cysts in the brain^43^ and reducing parasite burden in mesenteric lymph nodes^44^. Knowledge of very early *T. gondii* infection events in intestinal tissues is limited, and a possible contribution of FN-λ is unknown.

In the present study, we investigated the role of IFN-λ for restriction of *T. gondii* upon oral infection with tissue cysts. Our results demonstrate a significant impact of IFN-λ signaling to *T. gondii* control at the initial infection site. IFN-λ signaling in intestinal epithelial cells and neutrophils is thereby required to limit systemic spread of the parasite resulting in decreased burden of tissue cysts in the brain. IFN-λ also potentiated the *T. gondii*-specific humoral immune responses by enhancing the production of immunoglobulin IgG1. These are novel aspects of the infection biology of the parasite and might help to improve current and/or to develop new treatment strategies against toxoplasmosis.

## Results

### IFN-λ protects mice from lethal oral *T. gondii* infection

Because of the growing evidence that interferon lambda (IFN-λ) is protective against a variety of mucosal pathogens^2,9–13,19,20^, we examined the importance of IFN-λ signaling upon oral *T. gondii* infection (**Suppl. Figure 1)**. In IFN-λ receptor-deficient (*Ifnlr1^-/-^*) male mice, a significant increase in mortality compared to wild type mice was observed 10 days after oral administration of freshly prepared *T. gondii* ME49-derived tissue cysts (**Figure 1A**). Susceptibility of *Ifnlr1^-/-^* mice was reflected by increased weight loss compared with wild type (wt) animals (**Figure 1A)**. No apparent differences could be observed between *Ifnlr1^-/-^* and wt female mice infected with *T. gondii* ME49-derived tissue cysts in the same experiments. In these cases, all animals succumbed to infection (**Figure 1B**).

**Figure 1.**
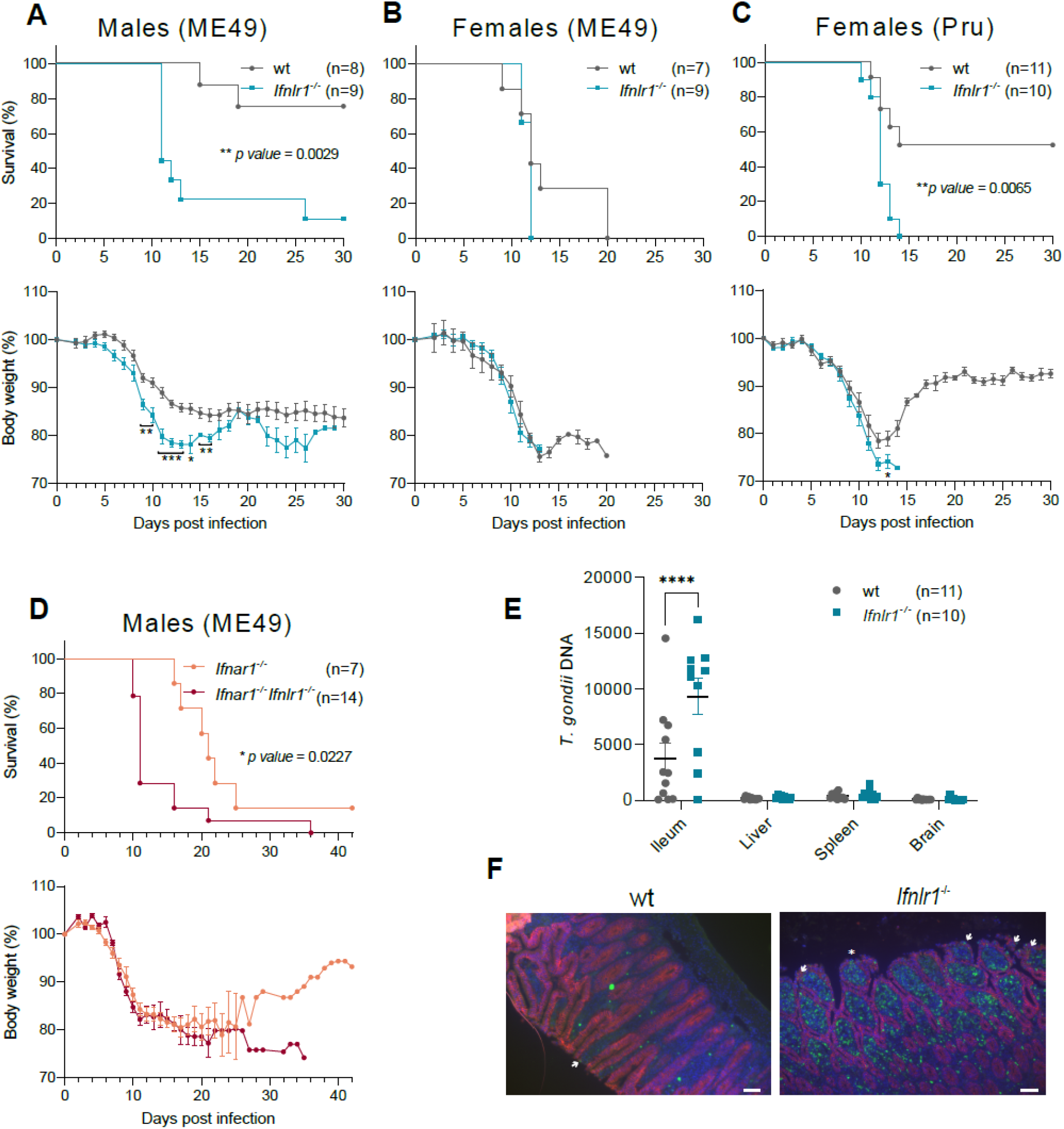
*Ifnlr1^-/-^* mice are highly susceptible to *T. gondii* oral infection. **A, B, D.** wt, *Ifnlr1^-/-^, Ifnar1^-/-^* or *Ifnar1^-/-^Ifnlr1^-/-^* mice were infected with 5 *T. gondii* ME49 or **C.** 10 *T. gondii* Pru-tdTomato tissue cysts. Weight loss and survival were monitored daily for 30 days. Data were pooled from two independent experiments. **A.** Survival (upper panel), \*\**p* = 0.0029 determined by Log-rank (Mantel-Cox) test; weight loss (lower panel), *p = 0.04, \*\**p* ≤ 0.002, \*\*\**p* ≤ 0.0007 determined by unpaired t test. **C.** Survival (upper panel), \*\**p* = 0.0065 determined by Log-rank (Mantel-Cox) test; weight loss (lower panel), \**p* = 0.03 determined by unpaired t test. **D**. Survival (upper panel), *\*p* = 0.0227 determined by Log-rank (Mantel-Cox) test. **E-F.** *T. gondii* replication in the intestine of *Ifnlr1^-/-^* and wt animals. *Ifnlr1^-/-^* and wt mice were infected with 15 *T. gondii* ME49-GFP-Luc tissue cysts for 9 days. **E.** *T. gondii* DNA (genomes) was quantified by qPCR. Data were pooled from two independent experiments, \*\*\*\**p* < 0.0001 determined by ANOVA with Tukey’s multiple-comparison test. **F**. *T. gondii* replication in the ileum of *Ifnlr1^-/-^* and wt mice from **E** was visualized by immunofluorescence. Arrows indicate infected IECs, the asterisk indicates damaged epithelium.

Any IFN-λ-dependent phenotype in females might have been masked by increased susceptibility to oral *T. gondii* infection due to reduced body weight compared with male mice. We therefore infected female mice with Pru-dtTomato-derived tissue cysts, a less virulent and cystogenic *T. gondii* strain^45^. *Ifnlr1^-/-^* female mice reached humane end points until 14 days post infection while a significant higher survival rate was observed in case of wt mice (**Figure 1C**). Susceptibility of *Ifnlr1^-/-^* mice was reflected by increased weight loss compared with wt animals (**Figure 1C**), demonstrating that the protective effect of IFN-λ upon oral *T. gondii* infection is not sex-dependent.

To assess if type I and III interferons have additive protective effects, we infected interferon alpha receptor-deficient (*Ifnar1^-/-^*) and double deficient *Ifnar1^-/-^Ifnlr1^-/-^* male mice with *T. gondii* ME49-derived tissue cysts. *Ifnar1^-/-^Ifnlr1^-/-^* mice (**Figure 1D**) were highly susceptible, closely resembling the phenotype of single deficient *Ifnlr1^-/-^* mice (**Figure 1A**). *Ifnar1^-/-^* mice were initially more resistant and started to show severe signs of disease only after day 16 post infection (**Figure 1D**). These results demonstrate that both types of interferon play a non-redundant role in host defense against *T. gondii.* While IFN type I is important during the chronic phase, as previously reported^43^, IFN-λ is rather required in the acute phase of *T. gondii* oral infection.

To evaluate if *Ifnlr1^-/-^* mice fail to inhibit *T. gondii* replication, we quantified parasite burden in different organs by qPCR 9 days post oral infection. *T. gondii* burden was significantly increased in the ileum of *Ifnlr1^-/-^* compared to wt mice (**Figure 1E**) but no differences were found in liver, spleen or brain (**Figure 1E**). Immunofluorescence analysis of ileum sections at day 9 post *T. gondii* oral infection confirmed increased parasite replication in the lamina propria and intestinal epithelial cells (IECs) of *Ifnlr1^-/-^* compared to wt mice (**Figure 1F**). Thus, IFN-λ is required for the control of *T. gondii* replication at the initial infection site.

### IFN-λ signaling in IECs and immune cells mediates protection against oral *T. gondii* infection

To dissect the impact of IFN-λ signaling in immune cells and IECs on *T. gondii* control, we generated bone marrow (BM) chimeric mice and infected them orally with *T. gondii* Pru-tdTomato-derived tissue cysts. A significant increased susceptibility of *Ifnlr1^-/-^* recipient mice that received BM of *Ifnlr1^-/-^* donor mice was observed compared to wt recipient mice that received wt BM (**Figure 2A**), hence reproducing our initial findings (**Figure 1A and C**). Interestingly, an intermediate phenotype was observed for either of the heterologous chimeras (*Ifnlr1^-/-^* recipient mice that received wt BM and wt recipient mice that received *Ifnlr1^-/-^* BM) (**Figure 2A**). These results suggest that protection of mice from lethal oral *T. gondii* infection requires IFN-λ signaling in both, hematopoietic stem cell- (HSC) and non-HSC-derived cells.

**Figure 2.**
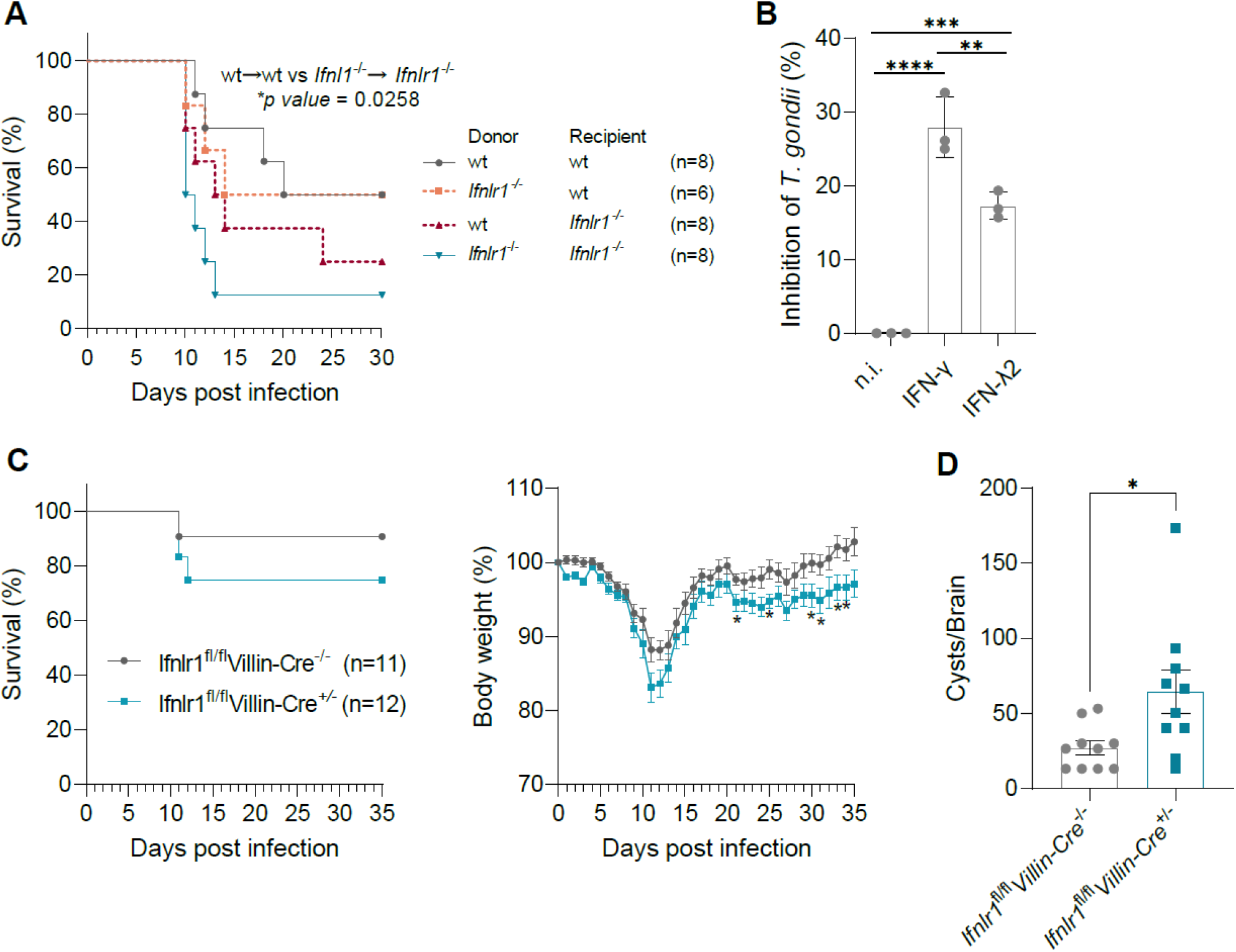
IECs and neutrophils contribute to IFN-λ-mediated protection from oral *T. gondii* infection. **A.** Bone marrow-chimeric mice were infected with 10 *T. gondii* Pru-tdTomato tissue cysts. Survival was monitored daily for 30 days. Survival, \**p* = 0.0258 determined by Log-rank (Mantel-Cox) test. **B.** *T. gondii* replication is inhibited in neutrophils. Neutrophils were primed for 8 h with indicated cytokines and infected with *T. gondii* ME49-GFP-Luc for 10 h. *T. gondii* inhibition was assessed by FACS. Results represent the mean and SEM of three independent experiments performed in duplicates or triplicates, **p = 0.0058, \*\*\**p* = 0.0004, \*\*\*\**p* <0.0001 determined by ANOVA with Tukey’s multiple-comparison test. **C-D.** The absence of IFNLR1 in the intestine leads to reduced weight recovery and higher cyst burden. **C, D.** *Ifnlr1*^fl/fl^*Villin-Cre*^+/-^ and *Ifnlr1*^fl/fl^*Villin-Cre*^-/-^ mice littermates were infected with 10 *T. gondii* Pru-tdTomato tissue cysts. Survival and weight loss were monitored daily for 35 days. Data were pooled from three independent experiments. **C.** Weight loss (right hand panel), \**p* ≤ 0.043 determined by Unpaired t test. **D.** Cyst burden in the brain was determined 35 days post infection, *p = 0.0238 determined by Unpaired t test.

Among other immune cell types, neutrophils are preferentially infected by *T. gondii* in the lamina propria^35^ and have been shown to exert different anti-*T. gondii* effector activities^46–48^. Furthermore, expression of IFNLRs has been demonstrated in murine neutrophils ^14,23,24^. Therefore, we investigated whether IFN-λ signaling in neutrophils inhibits *T. gondii* replication. Priming of mouse BM-derived neutrophils with 3 ng ml^-1^ of IFN-λ2 or IFN-γ resulted in saturated gene expression levels of *Isg15* and *Gbp2* (**Suppl. Figure 2**). Next, neutrophils were stimulated with 3 ng ml^-1^ of either IFN-γ or IFN-λ for 8 h and subsequently infected with *T. gondii* ME49-GFP-Luc. After 10 h of infection, *T. gondii* growth was determined by flow cytometry (**Suppl. Figure 3**). We observed that both, IFN-γ- and IFN-λ2-stimulated neutrophils, were able to significantly inhibit *T. gondii* replication (**Figure 2B**). These results demonstrate that neutrophils contribute to *T. gondii* inhibition upon IFN-γ and IFN-λ stimulation and might explain the intermediate phenotype observed in one (wt recipient mice that received *Ifnlr1^-/-^* BM) of the heterologous BM chimeras (**Figure 2A)**.

To verify the role of IFN-λ in IECs (**Figure 2A**), we infected mice lacking IFNLR1 specifically in the intestinal epithelium (*Ifnlr1*^fl/fl^*Villin-Cre*^+/-^ with *T. gondii* Pru-tdTomato-derived tissue cysts. Infected *Ifnlr1*^fl/fl^*Villin-Cre*^+/-^ mice showed increased weight loss at day 20 post infection (**Figure 2C**) as well as increased burden of *T. gondii* tissue cysts in the brain compared to *Ifnlr1*^fl/fl^*Villin-Cre*^-/-^ animals (**Figure 2D**). A similar picture emerged when *Ifnlr1*^fl/fl^*Villin-Cre*^+/-^ mice were infected with tissue cysts derived from *T. gondii* ME49 (**Suppl. Figure 4A-B**).

Altogether, our data demonstrate that IFN-λ signaling in IECs and neutrophils protects mice against lethal oral *T. gondii* infection by limiting parasite dissemination from the initial infection site to other organs including the brain.

### IFN-λ-dependent control of *T. gondii* in intestinal ODMs is mediated by IRG proteins

Intestinal organoids (“miniguts”) allow to investigate the early events after *T. gondii* infection *ex vivo*^49^. We established small intestine Organoid-Derived Monolayers (ODMs)^49^ (**Suppl. Figure 5A**) to evaluate the role of IFN-λ in IECs *in vitro.* We found that ODMs secrete IFN-λ2/3 into the supernatant 48 h post *T. gondii* ME49 infection (**Suppl. Figure 5B**). Priming of IECs with 30 to 60 ng ml^-1^ of IFN-λ2 resulted in saturated expression levels of the representative ISGs *Mx1* and *Irgb6* (**Suppl. Figure 6A).** To assess the impact of type I, II and III IFN on *T. gondii* growth inhibition, ODMs were therefore stimulated with 60 ng ml^-1^ IFN-α_B/D_, IFN-γ or IFN-λ2 for 24 h and luciferase activity was measured after 48 h of infection with a *T. gondii* luciferase reporter strain (ME49-GFP-Luc) (**Figure 3A**). Stimulation with IFN-γ or IFN-λ2 led to ~80 % and ~40 % inhibition of *T. gondii* growth respectively, whereas inhibition of *T. gondii* growth in IFN-α_B/D_-stimulated ODMs was hardly detectable (**Figure 3A**). The anti-parasitic activity mediated by IFN-λ2 but not IFN-γ was completely abolished in *Ifnlr1^-/-^* derived ODMs confirming specificity of *T. gondii* inhibition by IFN-λ2 (**Figure 3B**). These results demonstrate a differential impact of each type of IFN for *T. gondii* control in mouse intestinal ODMs.

**Figure 3.**
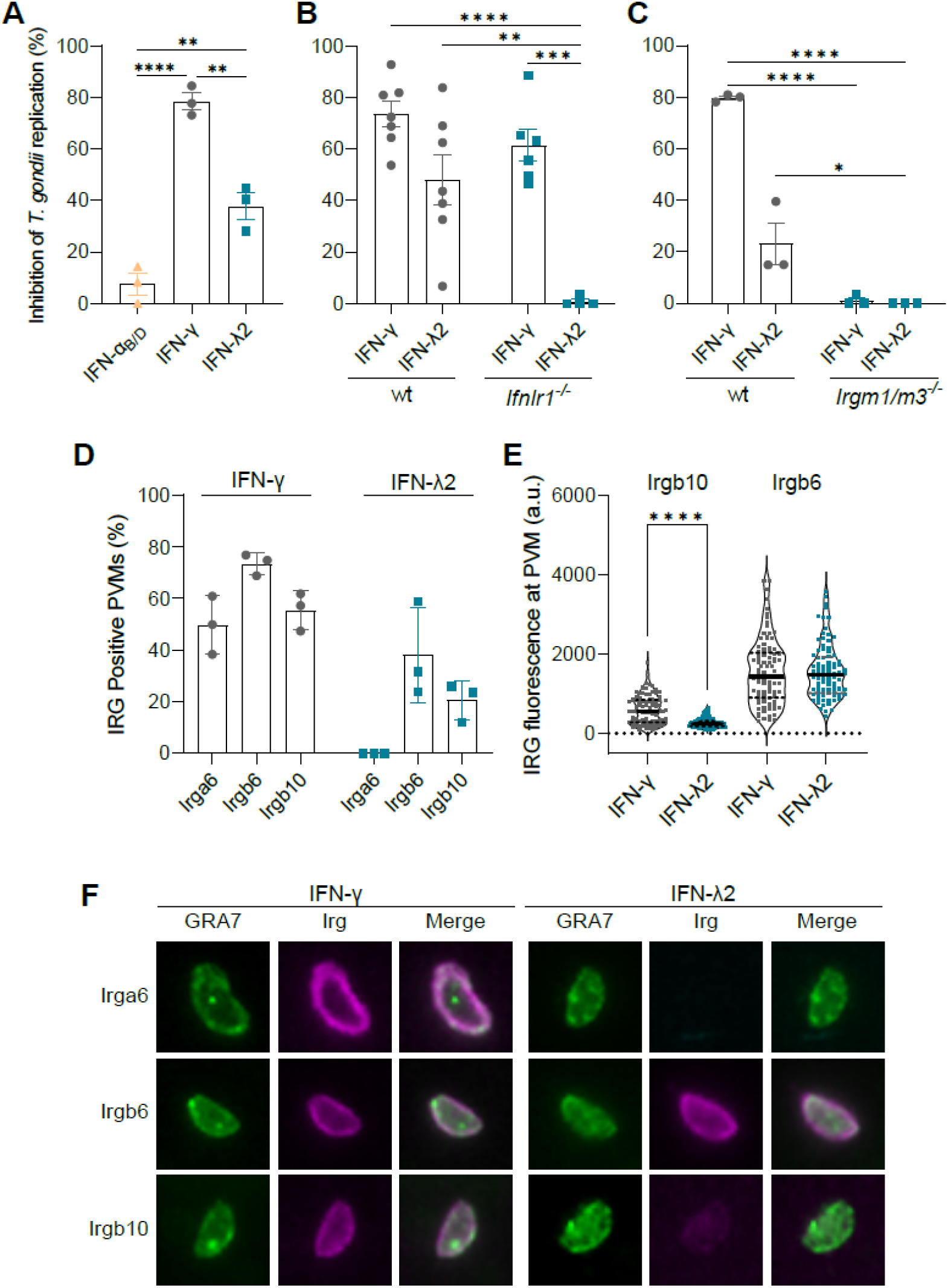
*T. gondii* replication is inhibited in ODMs. **A, B, C.** Organoid-Derived Monolayers (ODMs) were treated o/n with indicated cytokines and luciferase activity was measured 48 h post infection. **A.** *T. gondii* replication is inhibited by IFN-λ or IFN-γ in wt ODMs. Data represent the mean and SEM of three independent experiments performed in triplicates, \*\**p* ≤ 0. 0055, \*\*\*\**p* < 0.0001 determined by ANOVA with Tukey’s multiple-comparison test. **B.** *T. gondii* inhibition is lost in *Ifnlr1^-/-^* ODMs. Results represent the mean and SEM of 4-7 independent experiments performed in triplicates, \*\**p* = 0.0018, \*\*\**p* = 0.0002, \*\*\*\**p* < 0.0001 determined by ANOVA with Tukey’s multiple-comparison test. **C.** *T. gondii* inhibition is abrogated in *Irgm1/Irgm3^-/-^* ODMs. Data represent the mean and SEM of three independent experiments performed in duplicates or triplicates, \**p* = 0.0233, \*\*\*\**p* < 0.0001 determined by ANOVA with Tukey’s multiple-comparison test. **D-F.** IRG protein accumulation at the *T. gondii* PVM. ODMs were treated o/n with indicated cytokines and IRG proteins detected 2 h post *T. gondii* infection. **D.** Frequencies of IRG protein positive PVMs. 100 vacuoles were evaluated in each of three independent experiments. **E.** Intensities of IRG proteins at the PVM. 30 vacuoles were evaluated in three independent experiments respectively, \*\*\*\**p* < 0.0001 determined by Unpaired t test. **F.** Fluorescent images of IRG proteins at the PVM 2 h post infection.

An essential mechanism of *T. gondii* control in mice is constituted by the IFN-γ-inducible Immunity-Related GTPases (IRG proteins)^40^. Whereas effector IRG protein localization at the PVM is a prerequisite for membrane disintegration and parasite clearance^42,50–52^, regulator IRG proteins (Irgm1, Irgm2 and Irgm3) keep the effector IRG proteins in an inactive GDP-bound state at endomembranes in uninfected cells^53–55^. *Irgm1/Irgm3^-/-^* mice are highly susceptible to *T. gondii* infection due to mislocalisation of effector IRG proteins^54^. To evaluate the requirement of the IRG system for IFN-mediated growth inhibition of *T. gondii* in ODMs, we infected wt- and *Irgm1*/*Irgm3^-/-^*-derived ODMs after stimulation with IFN-γ or IFN-λ2 for 24 h with ME49-GFP-Luc and determined *T. gondii* growth inhibition at 48 h post infection. We found that *Irgm1/Irgm3^-/-^*-derived ODMs failed to inhibit *T. gondii* replication upon IFN-γ or IFN-λ2 stimulation (**Figure 3C**). These results demonstrate the importance of Irgm1/Irgm3-regulated IRG effector proteins for *T. gondii* control in intestinal ODMs.

To investigate the contribution of IRG proteins to *T. gondii* control at the initial infection site in more detail, we determined the expression levels of different *IRG* genes upon IFN-γ or IFN-λ2 treatment. While stimulation of ODMs with IFN-γ induced the expression of *Irga6, Irgb6, Irgb10* and *Irgd* but not *Mx1*, a classical IFN type I/III-inducible gene (**Suppl. Figure 6B**), IFN-λ2 treatment resulted in expression of *Irgb6, Irgb10, Irgd* and *Mx1* but not *Irga6* (**Suppl. Figure 6A-B**). Furthermore, we demonstrated the recruitment of Irgb6 and Irgb10 to the *T. gondii* PVM after IFN-λ2 treatment, although in lower frequencies compared to IFN-γ stimulation (**Figure 3D, F**). Whereas the mean fluorescent intensities of Irgb6 were essentially the same in IFN-γ- and IFN-λ2-stimulated ODMs, the mean fluorescent intensities of Irgb10 were higher in IFN-γ-compared with IFN-λ2-stimulated ODMs (**Figure 3E-F**). Whether the different patterns of expression of effector IRG proteins after stimulation with IFN-γ or IFN-λ2, that is partially reflected by accumulation at the *T. gondii*-derived PVM, can explain the differences observed in the magnitude of *T. gondii* inhibition (**Figure 3A, B, C**) still needs to be determined. Altogether, our data demonstrate that IFN-λ2 induces the expression and vacuolar accumulation of key anti-*T. gondii* proteins that are necessary to control parasite replication in the small intestine.

### IFN-λ treatment improves recovery and increases the specific *T. gondii* humoral response

Recombinant IFN-λ has been used as a therapeutic or prophylactic strategy to treat viral^12,23,28,56^ and *Cryptosporidium parvum* infections^17,18^. To evaluate the impact of IFN-λ treatment on oral *T. gondii* infections, we treated mice intraperitoneally with 1 ug ml^-1^ of IFN-λ1/3 daily from day −1 to day 7 of oral infection with tissue cysts of *T. gondii* Pru-dtTomato. No statistically significant differences in survival were observed between IFN-λ1/3-treated or PBS-treated control mice (**Figure 4A**). However, a significantly improved weight gain after day 17 post infection was observed in mice treated with IFN-λ1/3 in comparison to control mice (**Figure 4B**). Since reduced weight recovery after *T. gondii* infection correlated with enhanced cyst burden in the brain (**Figure 2C-D, Suppl. Figure 4A-B**), we determined the cyst numbers in the brain of IFN-λ1/3- and PBS-treated animals. *T. gondii* cyst counts were lower - although not significantly - in brains of mice treated with IFN-λ1/3 in comparison to PBS-treated control mice (**Figure 4C**) but no differences in cyst sizes were apparent between IFN-λ1/3-treated and PBS-treated mice (**Figure 4D**), indicating again that IFN-λ reduces *T. gondii* burden during the acute phase of infection rather than limiting the overall cyst growth during the chronic phase as it was reported for type I IFN^43^.

**Figure. 4.**
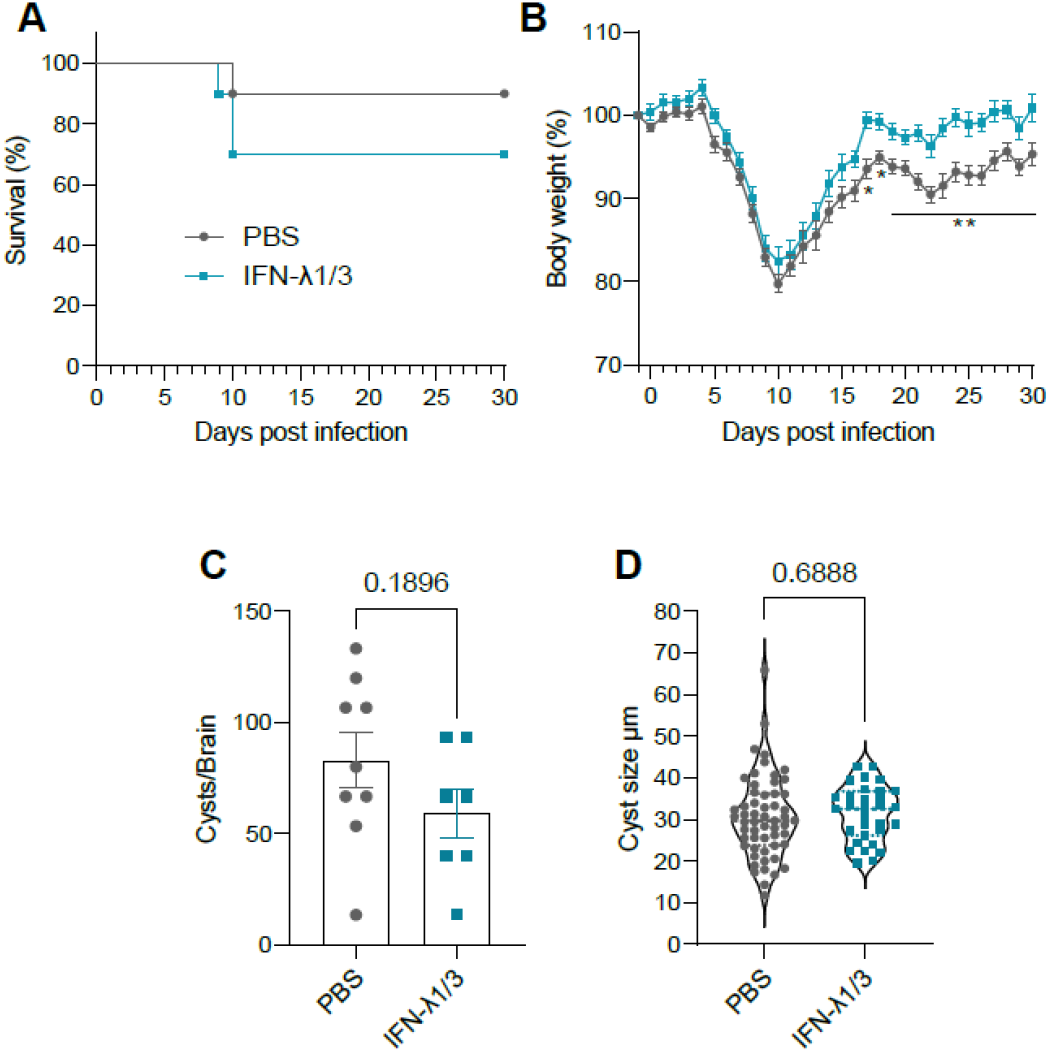
IFN-λ treatment improves recovery after oral *T. gondii* infection. **A-D.** Mice were treated (i.p. injection) with 1 μg of IFN-λ1/3 or mock-treated with PBS/0.1 % BSA from day −1 to day 7 of oral *T. gondii* infection with 10 Pru-tdTomato-derived tissue cysts and weight was monitored daily for 30 days. **A.** Survival. **B.** Weight loss, **\*\***p ≤ 0.0037 determined by Unpaired t test. **C, D.** Cyst numbers in the brain and cyst sizes were determined by DBA staining at 30 days post infection.

## Discussion

Coccidia are obligate intracellular parasites that can cause severe disease in humans and animals. Whereas most coccidian species have a narrow host tropism, *Toxoplasma gondii* (*T. gondii*) can infect almost all warm-blooded animals. In order to implement a control strategy against *T. gondii*, one main goal is to reduce the establishment of *T. gondii* tissue cysts and thereby limiting the risk of the parasite entering the human food chain. In the present study, we demonstrate the importance of interferon lambda (IFN-λ) signaling for *T. gondii* control at the initial site of infection, the intestine, consequently reducing the formation of tissue cysts (**Figure 5**).

**Figure 5.**
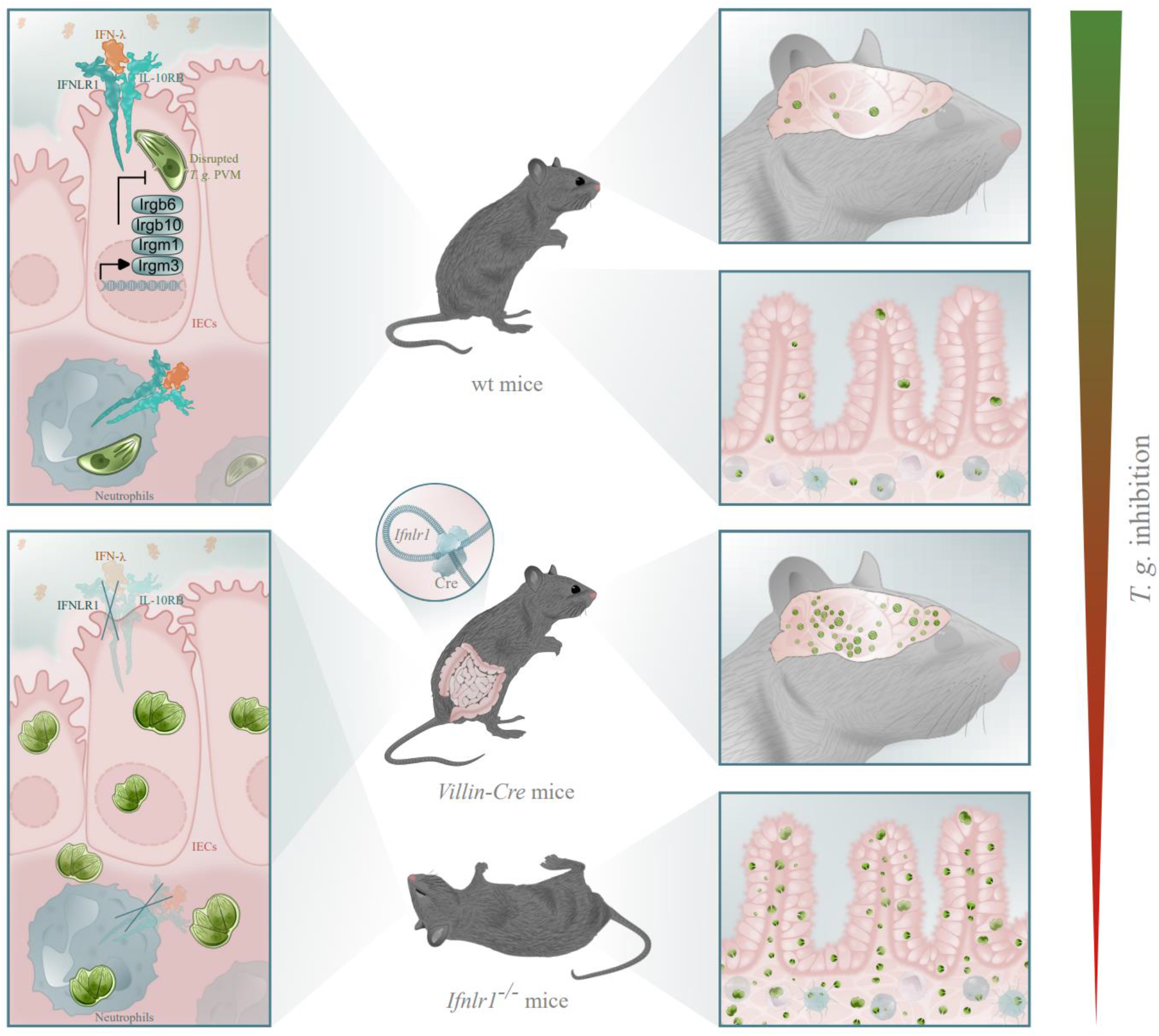
Importance of IFN-λ signaling for *T. gondii* control at the initial site of infection. Wild type (wt) mice survive oral *T. gondii* infection with tissue cysts. Loss of early IFN-λ-mediated *T. gondii* control in intestinal epithelial cells (IECs) (*Villin-Cre*) and neutrophils increases *T. gondii* replication in the intestine and colonisation to the brain, explaining the acute death of systemic interferon lambda receptor knockout (*Ifnlr1^-/-^*) mice. Whereas IRG proteins likely contribute to IFN-λ-mediated inhibition of *T. gondii* in IECs, the mechanism for parasite inhibition in neutrophils seems to be IRG-independent.

To pass the intestinal barrier, bradyzoites or sporozoites - contained in tissue cysts or oocysts, respectively - utilize different mechanisms^33,57^. Intestinal epithelial cells (IECs) are the most abundant epithelial cell type in the small intestine representing the first barrier that is invaded after release of bradyzoites or sporozoites from ingested tissue cysts or oocysts^58^. Infection of enterocytes at the apical side and release of tachyzoites after stage conversion and multiple rounds of intracellular replication at the basolateral side leads to systemic infection^33,34^. Therefore, intrinsic immune responses in enterocytes are important to limit parasite replication and dissemination^57,59,60^. Two families of IFN-γ-inducible GTPases (i.e. Immunity-Related GTPases (IRG proteins) and Guanylate Binding Proteins (GBP proteins)) are essential to control *T. gondii* infection in mice^40,41^, but their anti-*T. gondii* activities have never been evaluated in IECs. Three-dimensional multicellular organoids highly improve the reliability of host-pathogen interaction studies^61^. They are derived from stem cells or primary tissue and resemble the anatomy and physiology of intact organs. Organoid-Derived Monolayers (ODMs) possess most of the advantages of organoid structures and have been described previously as a suitable system to study *T. gondii* infection biology^49^. We found that stimulation of intestinal ODMs with IFN-γ induced the expression of IRG effector proteins (Irga6, Irgb6, Irgb10 and Irgd). Accumulation at the *T. gondii* parasitophorous vacuolar membrane (PVM) was concomitant with ~80 % inhibition of *T. gondii* replication (**Figure 3A-F**). IFN-λ stimulation resulted in somewhat lower expression levels of IRG effectors Irgb6, Irgb10 and Irgd, whereas Irga6 was not induced at all. Nevertheless, vacuolar accumulation of Irgb6 and Irgb10 could be detected and *T. gondii* replication was inhibited by IFN-λ up to 40 % (**Figure 3A-F**). Whether these differences in *T. gondii* control can be explained exclusively by the distinct expression patterns of IRG proteins upon IFN-λ or IFN-γ treatment and respective vacuolar IRG loading phenotypes awaits further investigation. However, in the absence of IRG regulator proteins Irgm1/Irgm3^54,55^ the anti-*T. gondii* effect mediated by both types of IFNs is completely abrogated (**Figure 3C**), demonstrating the importance of the IRG system for *T. gondii* control in IECs. At this point, we cannot rule out any contribution of GBP proteins to *T. gondii* control in IECs, especially because localization of Gbp1 and Gbp2 at the *T. gondii*-derived PVM is also regulated by Irgm1/Irgm3^54^.

We demonstrated that the specific deletion of the IFN-λ receptor (IFNLR) in IECs causes increased *T. gondii* colonization in the brain (**Figure 2D, Suppl. Figure 4B**). IFN-λ is produced after infection (**Suppl. Figure 5B**) by IECs and *T. gondii* replication is inhibited by IFN-λ in our *in vitro* system (**Figure 3A-C**). We therefore conclude that the IFN-λ-mediated immune response serves as an early host defense mechanism that limits *T. gondii* replication at the initial site of infection without provoking possible unfavourable immune responses mediated by IFN type I, similar to the distinctive role of IFN-λ in protecting mucosal surfaces during viral^2,5,62^ and *Cryptosporidium parvum (C. parvum*)^17,18^ infections. Future studies should determine the spacio-temporal details of the contribution of both IFNs during the early phases of infection. For example, by using a novel intestinal tissue microphysiological system in which the interaction between epithelium, endothelium and immune cells upon parasite infection can be analyzed^63^.

Systemic deletion of the IFNLR1 rendered mice highly susceptible to oral *T. gondii* infection (**Figure 1A, C, D**), however, conditional knockout (ko) of the IFNLR1 in IECs or bone marrow (BM) chimeric mice lacking the IFNLR1 in HSC-derived cells resulted in an intermediate phenotype (**Figure 2A, C**), indicating that IFN-λ signaling in both, epithelial and HSC-derived cells, contributes to protect against oral *T. gondii* infection. This is in contrast to experimental *C. parvum* infection of mice, where the IFN-λ-mediated antiparasitic activities were seemingly conferred exclusively by epithelial cells, even in immune-deficient *Rag2^-/-^Il2rg^-/-^* cells^18^. This is congruent with differences in the tissue tropism. While *C. parvum* infection is restricted to the intestine, *T. gondii* can establish a systemic infection, hence, systemic immune responses elicited against *T. gondii* are additionally required to limit dissemination of *T. gondii.*

Among other immune cells, neutrophils are preferentially infected by *T. gondii* in the lamina propria^35^ and express the highest IFNLR1 levels^22,27,64,65^. The weak induction of *IRG* and *GBP* genes in BM-derived neutrophils that we observed (**Suppl. Figure 8**) indicates that the IFN-λ2-mediated *T. gondii* control in neutrophils does not depend on IRG and GBP proteins and is mechanistically different from the parasite control elicited by IFN-λ in IECs.

IFN-λ acts on different immune cell types, thereby promoting or inhibiting different effector mechanisms. IFN-λ promotes ROS production by neutrophils to control *Aspergillus fumigatus* infection^65^, favours Th1 polarization through increased IL-12 production by dendritic cells^66^ or increases indirectly IFN-γ secretion by NK cells^67^. IFN-λ also acts as an immunomodulator in different inflammatory models (e.g. DSS-induced colitis and arthritis) on neutrophils by dampening ROS production, NET formation, degranulation and migration capacity, but maintaining phagocytic abilities. IFN-λ contributes to healing by maintaining the integrity and barrier function of epithelia at mucosal surfaces^22,24,64,68^. Mice orally infected with *T. gondii* develop enteritis due to the loss of Paneth cells, loss of barrier integrity and dysbiosis in an IFN-γ-dependent manner^60,69^. Whether IFN-λ promotes or modulates effector functions of other immune cells in addition to neutrophils during oral *T. gondii* infection still needs to be investigated.

Treatment of mice with IFN-λ augmented weight recovery and reduced *T. gondii* brain colonization without affecting *T. gondii* cysts sizes (Figure 4A-D), confirming our results that IFN-λ acts early during *T. gondii* infection by limiting parasite dissemination. IFN-λ serves as an mucosal adjuvant, promoting humoral responses in a thymic stromal lymphopoietin (TSLP)-dependent manner^2,29^. Interestingly, we found higher levels of secreted IgG1 after 30 days of *T. gondii* infection in the IFN-λ-treated group compared to non-treated mice (**Suppl. Figure 7B**). Our results are suggestive of a potential use of IFN-λ as an adjuvant for *T. gondii* vaccine development strategies, especially those that are delivered through mucosal surfaces.

Taken together, our work extends the repertoire of IFNs that contribute to the control of *T. gondii.* It advances our understanding of fundamental immunology against this worldwide zoonotic pathogen and might be relevant to enteric parasites *per se*.

## Methods

### *T. gondii* propagation

*T. gondii* tachyzoites ME49 (clone B7-21), ME49-GFP-Luc^1^ and Pru-Δ*hxgprt*-tdTomato (Pru-tdTomato)^2^ were propagated in confluent human foreskin fibroblasts (HFF1) in DMEM medium containing high glucose supplemented with 100 U ml^-1^ penicillin, 100 mg ml^-1^ streptomycin and 2 % fetal bovine serum.

### *T. gondii* tissue cysts preparation

For *T. gondii* tissue cyst preparation, C57BL/6 (BL/6) mice were infected with 200 to 500 *T. gondii* tachyzoites via intraperitoneal (i.p.) injection or with 5 to 15 tissue cysts freshly prepared from the brain of infected donor animals via oral gavage. Four to six weeks post infection, mice were euthanized by cervical dislocation. Brains were harvested and suspended in 2 ml PBS before mincing using a 18G and 20G needle respectively. Cyst numbers were determined via DBA-FITC staining at 20x magnification as described previously^3,4^.

### Animal strains and infection conditions

BL/6 mice were purchased from Janvier laboratories. B6.A2G-Mx1 mice carrying intact *Mx1* alleles (designated wt), congenic *B6.A2G-Mx1-Ifnar1^-/-^* mice lacking functional IFN-α receptors (designated *Ifnar1^-/-^*) and *B6.A2G-Mx1-Ifnlr1^-/-^* mice lacking functional IFN-λ receptors (designated *Ifnlr1^-/-^*) or double receptor-decient mice *B6.A2G-Mx1-Ifnar1^-/-^Ifnlr1^-/-^* (designated *Ifnar1^-/-^Ifnlr1^-/-^*) were described before^5^. B6.A2G-Mx1-*Ifnlr1^fl/fl^Villin-Cre*^+/-^ mice lacking functional IFN-λ receptors (IFNLR1) in intestinal epithelial cells (IECs) (designated *Ifnlr1*^fl/fl^*Villin-Cre*^+/-^) and control littermates B6.A2G-Mx1-*Ifnlr1*^fl/fl^*Villin-Cre*^-/-^ (designated *Ifnlr1*^fl/fl^*Villin-Cre*^-/-^) were described before^6^. Animals in all experimental groups were sex- and age-matched.

For survival experiments, mice were infected by oral gavage with 5 to 10 freshly prepared tissue cysts in a total volume of 200 μl sterile PBS. Infected mice were monitored daily for 30 to 35 days. Relative weight loss was calculated based on the weight at the day of infection.

For IFN-λ treatment, mice were treated i.p. with 1 μg of human IFN-λ1/3^7^ (proven to be cross-reactive in mice) or mock-treated with PBS/0.1 % BSA from day −1 to day 7 of oral infection with 10 *T. gondii* Pru-tdTomato-derived tissue cysts and monitored daily for 30 days.

Mice were kept under specific-pathogen-free conditions in the local animal facility (Department for Microbiology, Virology and Hygiene, Freiburg). All animal experiments were performed in accordance with the guidelines of the German animal protection law and the Federation for Laboratory Animal Science Associations. Experiments were approved by the state of Baden-Württemberg (Regierungspräsidium Freiburg; reference number G-19/89, G-20/155, and G-22/068).

### Generation of bone marrow chimeras

Five days before bone marrow transplantation (day −5 to day −1), wt and *Ifnlr1^-/-^* sex- and age-matched recipient mice received daily 20 mg of busulfan per kg body weight i.p. as previously described^8^. A suspension of bone marrow cells prepared from wt or *Ifnlr1^-/-^* donor mice were adoptively transferred into recipient animals. The recipient mice were given drinking water containing 2 g/l neomycin for 3 weeks after busulfan treatment. Eight weeks post BM transplantation, mice were orally infected with 5 *T. gondii* Pru-tdTomato-derived tissue cysts and monitored daily for 30 days.

### Isolation of bone marrow-derived neutrophils

Following the manufacturer’s protocol, neutrophils were negatively enriched using the EasySep™ Mouse Neutrophil Enrichment Kit (StemCell Technologies) from the bone marrow of adult wt mice. The cells were resuspended in PBS supplemented with 2 % FCS and 1 mM EDTA and cultured in RPMI medium supplemented with 100 U ml^-1^penicillin, 100 μg ml^-1^streptomycin, 10 % FCS, 1 mM sodium pyruvate (Capricorn Scientific) and 4 mM glutaMAX (Thermo Fisher Scientific).

### Generation of intestine Organoid-Derived Monolayers

Small intestine organoids from wt, *Ifnlr1^-/-^* and *Irgm1/Irgm3^-/-^* mice were generated according to StemCell technologies protocols. Stem cell enriched spheroids were cultured in Stem Cell enrichment medium (SC medium) as described before^9^.

To grow organoids as Organoid-Derived Monolayers (ODMs), 96 well plates, Ibidi μ-chambers or transwells were coated with 50 μl/well basement membranes (BME) diluted 1:20 in adDMEM^+/+/+^ o/n at 4°C or for at least 30 min at 37°C. The coating solution was aspirated and the cell culture kept at 37°C for another 30 min. Three to four days old organoids were recovered as described above, centrifuged at 300 g and 4°C for 5 min, and the pellets resuspended in 1 ml pre-warmed TrypLE (Thermo Scientific) + 10 μM Y-27632 (MedChemExpress). After incubation in a 37°C water bath for 2 min, the suspension was aspirated twice with a 1 ml syringe and a 20G needle pre-coated with organoid washing medium containing trypsin to create a single-cell suspension. To stop trypsinization, 10 ml adDMEM^+/+/+^ were added, the suspension centrifuged at 300 g and 4°C for 5 min, the pellet resuspended in an appropriate volume of ODM seeding medium, and the cell concentration determined using a hemocytometer (~ 6×10^4^ cells/cm^2^ were seeded). One day after seeding, the medium was exchanged to 90 % ODM differentiation medium. Additional medium changes were done every second day. ODMs were cultured for 6 days before IFN stimulation.

### *T. gondii* replication assays

Enriched bone marrow-derived neutrophils were primed with 3 ng ml^-1^of IFN-λ2^10^, IFN-γ (Peprotech) for 8 h and infected with *T. gondii* GFP-Luc at a multiplicity of infection (MOI) of 2. At 10 h post infection, neutrophils were recovered from culture plates using 200 μl of Accutase Cell Dissociation Solution (Sigma-Aldrich) for 25 min at 37°C. Cells were incubated in 200 μl FACS buffer containing 1 μl Zombie NIR fixable dye for 30 min to determine cell viability (Zombie Green™ Fixable Viability Kit, BioLegend). Cells were fixed for 15 min in 200 μl PFA 2 %. To avoid unspecific antibody binding, Fc blocking was performed using anti-FcyIII/II CD16/32 receptor antibody (Clone 93) for 10 min on ice. Cells were stained with fluorochrome-conjugated antibodies against cell surface markers for 20 min on ice (Table 1). Cells were finally washed in 200 μl of FACS buffer and resuspended in 300 μl FACS buffer. A FACS Canto II flow cytometer (Becton Dickinson) was used to collect 100.000 events and data were analyzed with FlowJo software.

**Table 1.** Antibodies.

| Species | Antigen | Type | Conjugat | Dilution | Reference / Origin |
| --- | --- | --- | --- | --- | --- |
| **Immunofluorescence primary antibodies** |  |  |  |  |  |
| mouse | Irga6 | moAb | - | 1:2000 | 10D7<sup>15</sup> |
| mouse | Irgb6 | moAb | - | 1:3000 | B34<sup>16</sup> |
| rabbit | Irgb10 | antiserum | - | 1:6000 | 940/6<sup>17</sup> |
| rabbit | E-Cadherin (24E10) | moAb | - | 1:400 | 3195 (Cell Signalling) |
| **Immunofluorescence secondary antibodies** |  |  |  |  |  |
| donkey | mouse IgG | poAb | Alexa 555 | 1:5000 | A31570 (Life Technologies) |
| donkey | rabbit IgG | poAb | Alexa 555 | 1:5000 | A31572 (Life Technologies) |
| donkey | rat IgG | poAb | Alexa 488 | 1:5000 | A21208 (Life Technologies) |
| **Flow cytometry antibodies** |  |  |  |  |  |
| rat IgG2a, κ | anti-mouse Ly-6G (1A8) | moAb | APC | 0.06 µg per 10<sup>6</sup> cells | 127613 (Biolegend) |
| rat IgG2b, κ | anti-mouse CD45 (30-F11) | moAb | PerCP | 0.25 µg per 10<sup>6</sup> cells | 103129 (Biolegend) |
| rat IgG2b, κ | purified anti-mouse CD16/32 (93) | moAb | - | 1.0 µg per 10<sup>6</sup> cells | 101301 (Biolegend) |

Percent of *T. gondii* infected neutrophils (CD45^+^, Ly6G^+^, GFP^+^) was compared between treated and non-treated conditions. Percent of *T. gondii* inhibition was defined as “*i* = 100 - [% of (CD45^+^, Ly6G^+^, GFP^+^) tretated] / [% of (CD45^+^, Ly6G^+^, GFP^+^) untretated] * 100”.

To evaluate inhibition of *T. gondii* replication by IFNs in IECs, ODMs were primed o/n with 60 ng ml^-1^ IFN-λ2^10^, IFN-γ (Peprotech) or IFN -α_B/D_^11^ and infected with *T. gondii* ME49-GFP-Luc at a MOI of 0.25 for 48 h. Cells were washed once with PBS and lysed for at least 1 h with 40 μl 1x passive lysis buffer (Promega) at RT. 20 μl of lysate were transferred to a white flat-bottom 96-well plate (Thermo Scientific) and luciferase activity was measured in a Tecan infinite 200Pro by automatic injection of 50 μl luciferase assay substrate (Promega) and 10 sec integration time.

Inhibition of *T. gondii* replication was calculated as “*i* = 100% - (*L*_treated_ / *L*_mock_) * 100%”, where “*i*” is the inhibition of *T. gondii* replication and “*L*” is the luminescence in the IFN-treated or untreated wells. Negative *T. gondii* inhibition values were set to zero.

### *T. gondii* quantification and *ISG* expression by qPCR

Wt and *Ifnlr1^-/-^* mice were infected orally with 15 *T. gondii* ME49-GFP-Luc-derived tissue cysts in a total volume of 200 μl sterile PBS. After 9 days of infection, biopsies from ileum, spleen, liver and brain were taken and preserved in DNA/RNA shield (Zymo Research) at −80°C until DNA/RNA isolation. RNA and DNA was isolated with the Direct-zol^TM^ DNA/RNA kit (Zymo Research). Parasite load was quantified from purified DNA by a probe-based qPCR using specific primers that amplify the 529 bp repetitive element (RE) in the parasite genome (Table 2)^12^. Purified *T. gondii* DNA was used to create a standard curve for calculation of parasite load. DNA was amplified using Luna® Universal Probe qPCR Master Mix (NEB England Biolabs).

**Table 2.**
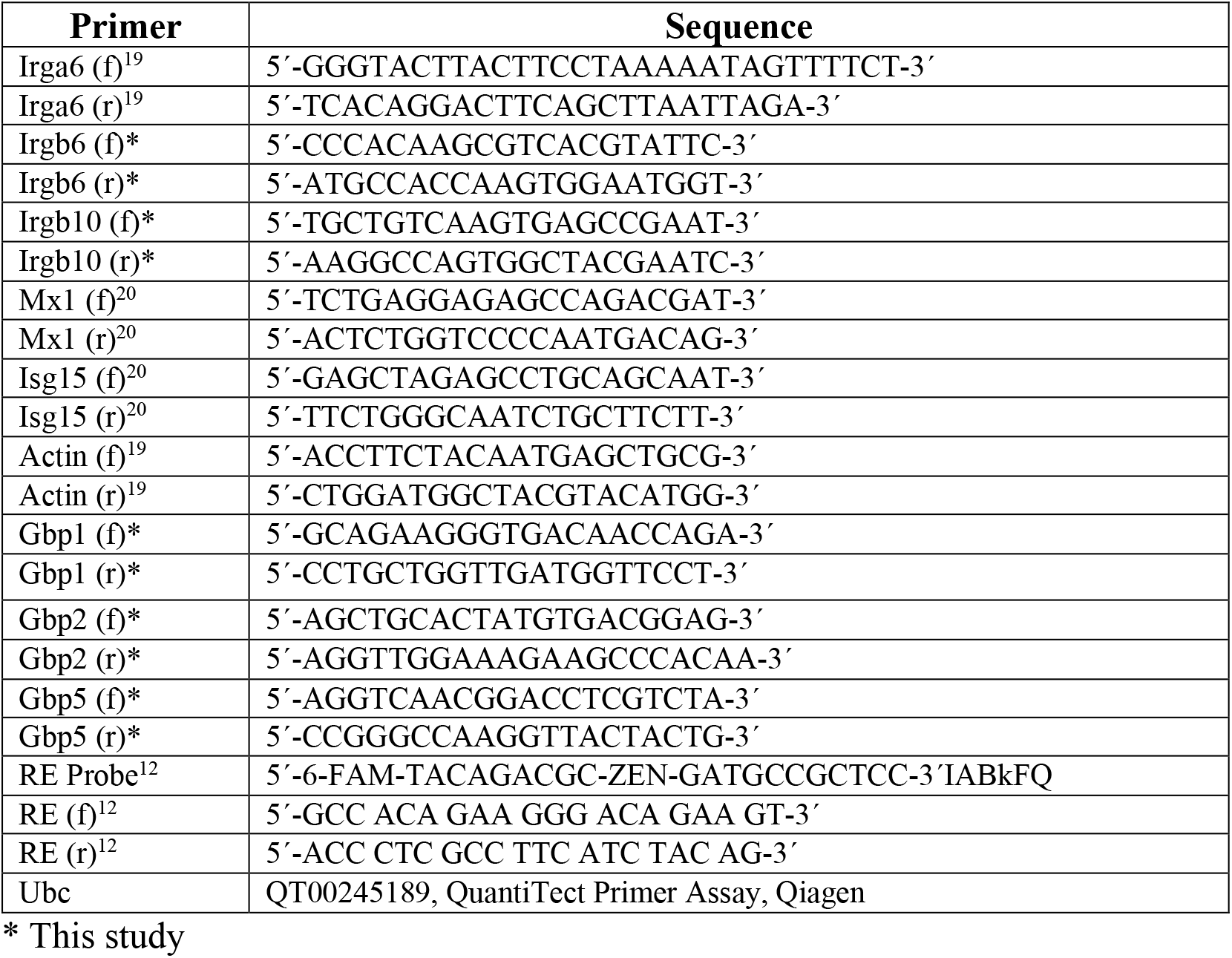
Primers.

For *ISG* induction, purified neutrophils or ODMs were primed with different concentrations of IFN-γ or INF-λ2 for 4 h. Afterwards, RNA was purified using the Direct-zol™ RNA Miniprep Kit (Zymo Research) according to the manufacturer’s protocol. Complementary DNA (cDNA) was generated for each replicate using the LunaScript RT Supermix (New England Biolabs) based on the manufacturer’s instructions. The cDNA served as template for the amplification of genes of interest (Table 2), using SYBR green I containing Luna® Universal qPCR Master Mix (NEB England Biolabs). The qPCR was performed using the QuantStudio 5 Real-Time PCR System (Applied Biosystems by Thermo Fisher Scientific). The increase in mRNA expression was determined by the 2-ΔΔCt method relative to the expression of the house-keeping gene *Ubc* or ΔCt relative to *Actin*.

### ELISA

*T. gondii*-specific antibodies in serum of IFN-λ1/3 treated mice were determined by ELISA as described previously^3^. Briefly, high-binding 96-well microtiter plates (MaxiSorp, Nunc) were coated with total *T. gondii* antigen and incubated overnight at 4°C. Next, the ELISA plates were washed four times with washing buffer (PBS containing 0.05 % Tween 20) and blocked with 1 % BSA in PBS for 1 h at 37°C. Plates were washed four times with washing buffer. Afterwards, 1:128 diluted serum were added and incubated for 1 h at room temperature. Plates were washed four times with washing buffer. Horseradish peroxidase-labelled antibodies directed against either total IgG (62-6520, Invitrogen) or IgG1 (A10551, Invitrogen) were added to each well and incubated for 1 h at room temperature. Plates were washed four times and incubated with tetramethylbenzidine (TMB) substrate (Biologend) for 10 min at room temperature. The reaction was stopped by adding 0.5 M H2SO4 and the absorbance was measured at 450 nm and 570 nm (background). *T. gondii* specific IgG or IgG1 values were calculated relative to values from non-infected mice.

IFN-λ2/3 concentration was determined by commercial sandwich ELISA (R&D Systems) from supernatants infected or not with *T. gondii* after 24 or 48 h post infection.

### Immunofluorescence

Antigen retrieval in deparaffinized paraformaldehyde-fixed ileum tissue sections from wt and *Ifnlr1^-/-^* mice was performed with 0.01 M sodium citrate buffer as previously described^13^. Slides were blocked with 10 % normal donkey serum (Jackson ImmunoResearch) and stained o/n with rat anti GRA7 (*T. gondii* PVM marker) and E-Cadherin (Cell Signalling) followed by 1 h incubation with the appropriate Cy3-, or Cy5-conjugated secondary antibodies and DAPI. Slides were mounted in Fluor Save Reagent (Calbiochem). Tissue sections were visualized using a Zeiss Axioplan 2 non-inverted fluorescence microscope.

ODMs were infected with *T. gondii* ME49 for 2 h at MOI 4. Monolayers were washed two times with PBS and fixed for 30 min at RT with 4 % PFA. Cells were permeabilized and stained as previously described^14^. Antibodies and dilutions are listed in Table 1. Intensities were determined by taking the average of 4 intensity values along 2 lines crossing the PV perpendicularly subtracted by the respective background fluorescence, as described previously^80^. The measurements were done using the Fiji/ImageJ software with a custom macro (code can be found at https://github.com/Kartoffelecke/PVM-profiler). Pictures were taken on the Zeiss Observer 7 with a 40x magnification.

### Statistical analysis

All statistical analyses were performed using GraphPad Prism 9.1 software. P-values were determined by an appropriate statistical test. One-way ANOVA followed by Tukey’s multiple comparison was used to test differences between three or more groups. Depending on the data distribution, Student’s t-test or Mann Whitney test was used for two-group comparisons. For *in vivo* experiments, a log-rank Mantel-Cox test was used to test survival differences between groups. All error bars indicate the mean and standard error of the mean (SEM) of at least three independent experiments. P-values; ****p < 0.0001, ***p < 0.001, **p < 0.0, *p < 0.05, n.s. no significant.

### Intestinal organoid media^9^

#### adDMEM^+/+/+^

adDMEM/F-12 (Gibco 12634028)
+ 75 U mL^-1^ penicillin
+ 75 μg mL^-1^ streptomycin (Gibco 15140122)
+ 10 mM HEPES pH 7.5
+ 1x GlutaMax (Gibco 35050061)

### Stem Cell enrichment (SC) organoid medium

adDMEM/F-12
+ 50 % L-WRN conditioned medium
+ 20 % R-spondin conditioned medium
+ 10 % noggin conditioned medium
+ 75 U mL^-1^ penicillin + 75 μg mL^-1^ streptomycin (Gibco 15140122)
+ 10 mM HEPES pH 7.5
+ 1x GlutaMax (Gibco 35050061)
+ 1 mM N-acetylcystein (Sigma)
+ 10 mM nicotinamid (Sigma)
+ 1x B27 supplement (Gibco 17504044)
+ 1x N2 supplement (Gibco 17502048)
+ 50 ng mL^-1^ mEGF (StemCell Technologies)
+ 500 nM A83-01 (StemCell Technologies)
+ 10 μM SB202190 (StemCell Technologies)

### Organoid washing medium

adDMEM^+/+/+^ + 10 % FCS

### Organoid freezing medium

adDMEM^+/+/+^ + 20 % FCS + 10 % DMSO

### ODM seeding medium

adDMEM/F-12
+ 50 % L-WRN conditioned medium
+ 20 % R-spondin conditioned medium
+ 10 % noggin conditioned medium
+ 75 U mL^-1^ penicillin + 75 μg mL^-1^ streptomycin (Gibco 15140122)
+ 10 mM HEPES pH 7.5
+ 1x GlutaMax (Gibco 35050061)
+ 1 mM N-acetylcystein (Sigma)
+ 10 mM nicotinamid (Sigma)
+ 1x B27 supplement (Gibco 17504044)
+ 1x N2 supplement (Gibco 17502048)
+ 50 ng/mL mEGF (StemCell Technologies)
+ 10 μM Y-27632

### ODM differentiation medium

adDMEM/F-12
+ 20 % R-spondin conditioned medium
+ 10 % noggin conditioned medium
+ 75 U mL^-1^ penicillin + 75 μg mL^-1^ streptomycin (Gibco 15140122)
+ 10 mM HEPES pH 7.5
+ 1x GlutaMax (Gibco 35050061)
+ 1 mM N-acetylcystein (Sigma)
+ 10 mM nicotinamid (Sigma)
+ 1x B27 supplement (Gibco 17504044)
+ 1x N2 supplement (Gibco 17502048)
+ 50 ng mL^-1^ mEGF (StemCell Technologies)

## Acknowledgments

We are especially thankful for all support from the Institute of Virology under the direction of Hartmut Hengel. We thank Philipp P. Petric, Precious Cramer and Peter Reuther from the University Medical Center for technical assistance. We are equally grateful to Aladin Haimovici from the University Medical Center Freiburg for providing organoid conditional media and for helpful discussions. We thank Frank Seeber and Christian Klotz from the Robert Koch-Institute very much for assistance with generation and maintenance of organoids. Jonathan C. Howard from the Fundação Calouste Gulbenkian generously provided antibodies against IRG proteins. Rune Hartmann and Hans Hendrik Gaad from the Department of Molecular Biology and Genetics, Aarhus University, kindly provided recombinant IFN-λ2. T.S. received funding from the Deutsche Forschungsgemeinschaft STE 2348/2-1 and STE 2348/2-2, and University of Freiburg, Medical Faculty, Research Commission STE 2134/20. M.M.L. received funding (Research Grants–Doctoral Programmes in Germany) from the German Academic Exchange Service (DAAD). G.T. received funding from the US National Institutes of Health (AI145929 and AI148243). The funders had no role in study design, data collection and interpretation, or the decision to submit the work for publication.

## Author contributions

Mateo Murillo-León: Conceptualisation; Formal analysis; Investigation; Methodology; Writing-original draft; Writing-review & editing. Aura Maria Bastidas Quintero: Formal analysis; Investigation; Methodology; Writing-review & editing; Niklas Endres: Formal analysis; Investigation; Methodology; Writing-review & editing. Daniel Schnepf: Formal analysis; Investigation; Methodology; Writing-review & editing. Estefanía Delgado-Betacourt: Methodology; Writing-review & editing. Annette Ohnemus: Investigation; Methodology; Writing-review & editing Gregory Alan Taylor: Resources; Writing-review & editing. Martin Schwemmle: Resources; Writing-review & editing. Peter Staeheli: Formal analyisis, Investigation; Methodology; Writing-review & editing. Tobias Steinfeldt: Conceptualisation; Resources; Data curation; Validation; Supervision; Fundig acquisition; Project Administration; Formal analysis; Methodology; Writing-original draft; Writing-review & editing.

## Data availability

This study includes no data deposited in external repositories.

