## Supplementary Information for "IFN-λ is protective against lethal oral *Toxoplasma gondii* infection"

### Supplementary Figure legends

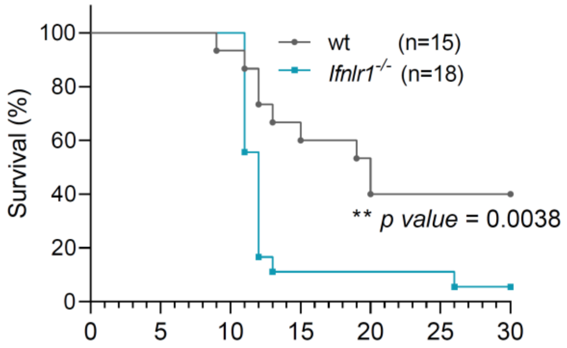


**Supplementary Figure 1. *Ifnlr1*$^{-/-}$ mice are highly susceptible to *T. gondii* oral infection.** Wt and *Ifnlr1*$^{-/-}$ mice were infected with 5 *T. gondii* ME49 tissue cysts. Survival was monitored daily for 30 days. Data were pooled from two independent experiments, $^{**}p$ = 0.0038 determined by Log-rank (Mantel-Cox) test.

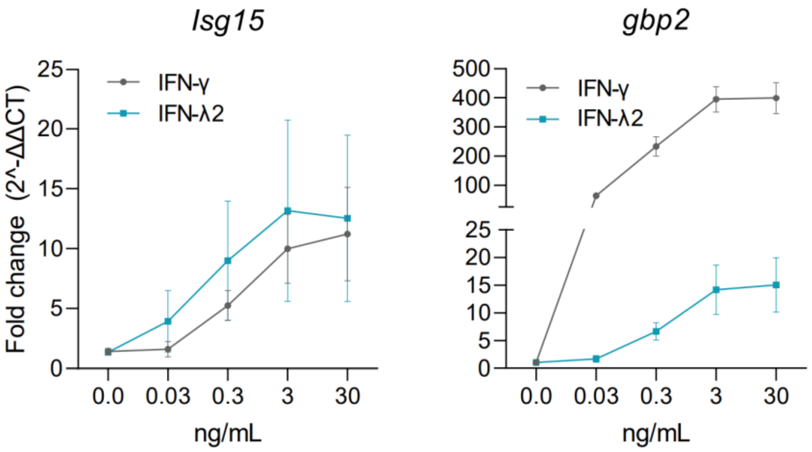


**Supplementary Figure 2. Titration of IFN-γ and IFN-λ2.** Neutrophils from the bone marrow of wt mice were stimulated for 4 h with increasing concentrations of IFN-γ or IFN-λ2 [0.003, 0.03, 0.3, 3 and 30 ng $ml^{-1}$]. Expression of *Isg15* and *Gbp2* was quantified relative to *Ubc*. Symbols represent means ± SD. Data correspond to the mean of four independent experiments performed in triplicates.

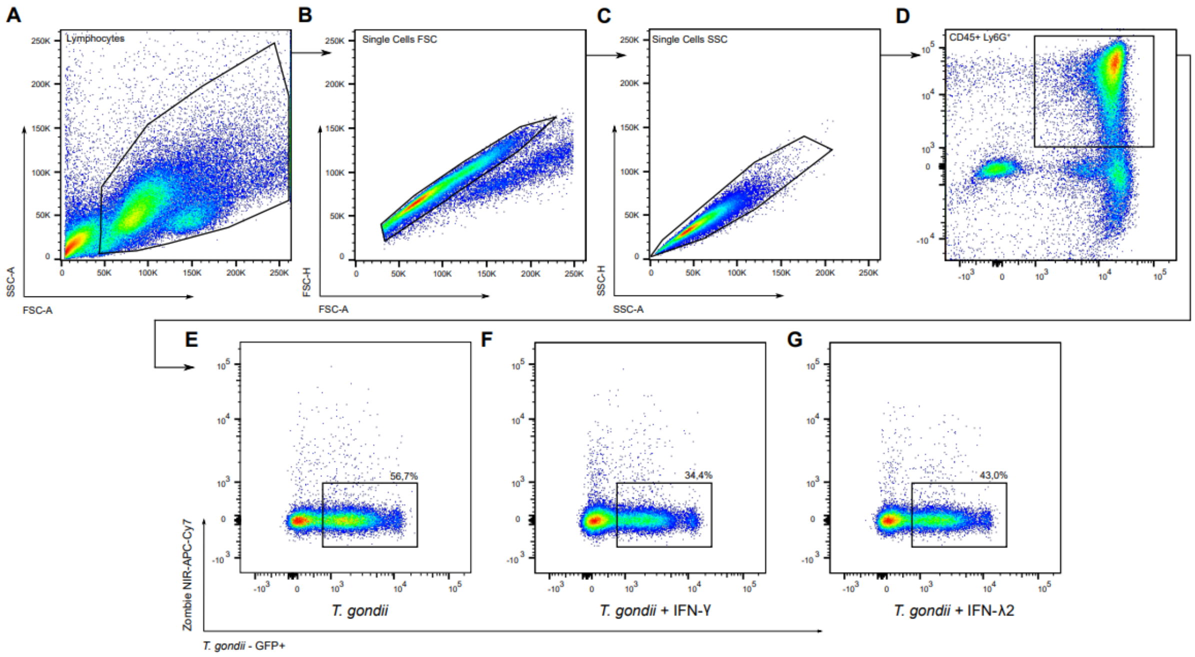


**Supplementary Figure 3**. **FACS gating strategy.** Cells were negatively isolated from the bone marrow of wt mice. Cells were infected with *T. gondii* ME49-GFP-Luc at MOI 2 for 10 h after priming with 3 ng $ml^{-1}$IFN-γ or IFN-λ2 for 8 h. Following FSC-SSC (a), singlet gating (b, c) and leukocyte neutrophil gating (CD45+ Ly6G+) (d), the proportion of GFP+ live cells were defined as infected (E-G).

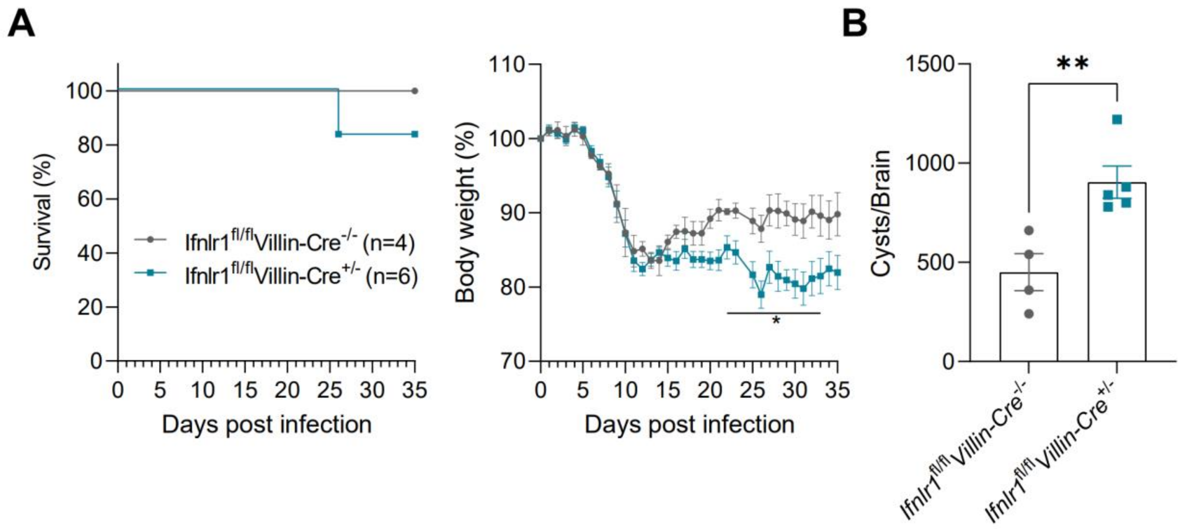


**Supplementary Figure 4. The absence of IFNLR1 in the intestine leads to reduced weight recovery and higher parasite cyst burden after oral *T. gondi* infection. A, B.** *Ifnlr1*$^{fl/fl}$*Villin-Cre*$^{+/-}$ and *Ifnlr1*$^{fl/fl}$*Villin-Cre*$^{-/-}$ mice littermates were orally infected with 10 *T. gondii* ME49-derived tissue cysts. Survival and weight loss of the animals were monitored daily for 35 days. Data represent one single experiment. **A.** Weight loss (right hand panel), *$p \leq 0.043$ determined by Unpaired t test. **B.** Cyst burden in the brain was determined by DBA staining at 35 days post infection, **$p = 0.0077$ determined by Unpaired t test.

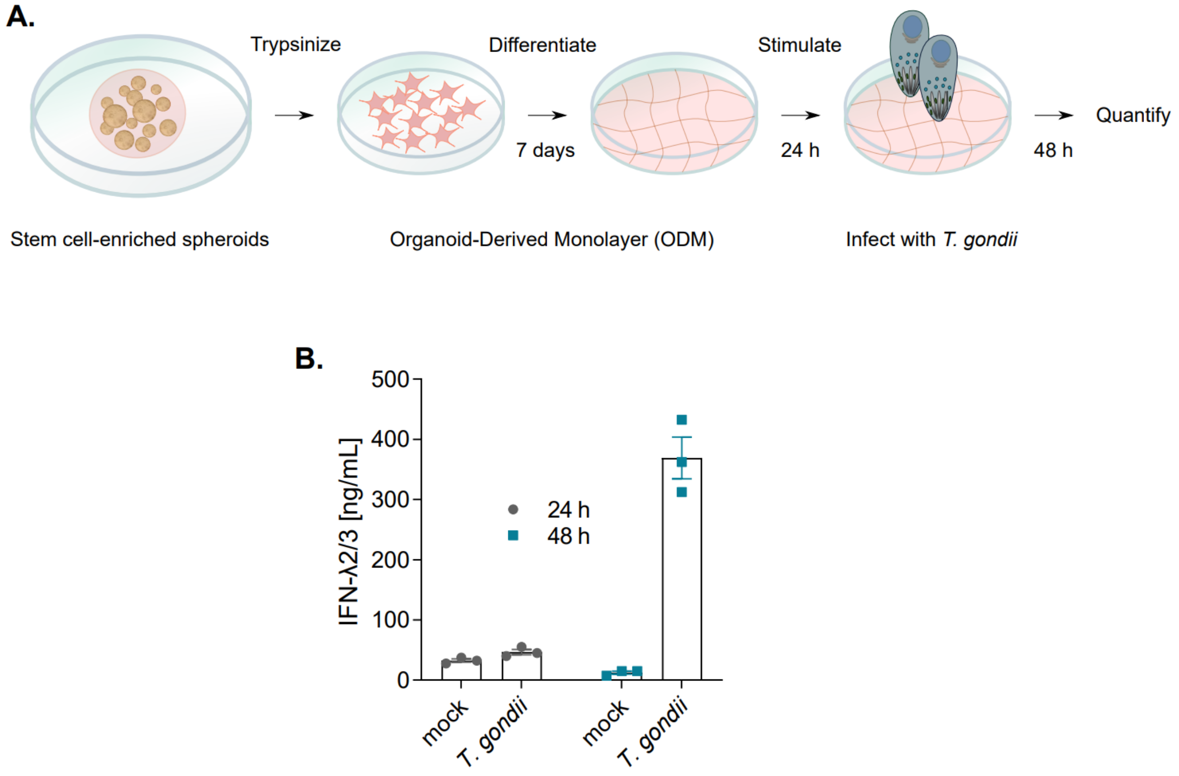


**Supplementary Figure 5**. **A.** Schematic representation of mouse intestine ODM establishment and *T. gondii* infection. **B.** IFN-λ2/3 is secreted after *T. gondii* infection of intestine-derived ODMs. Mouse small intestine ODMs were infected with *T. gondii* ME49-GFP-Luc for 24 h or 48 h and IFN-λ2/3 was detected in supernatants by ELISA.

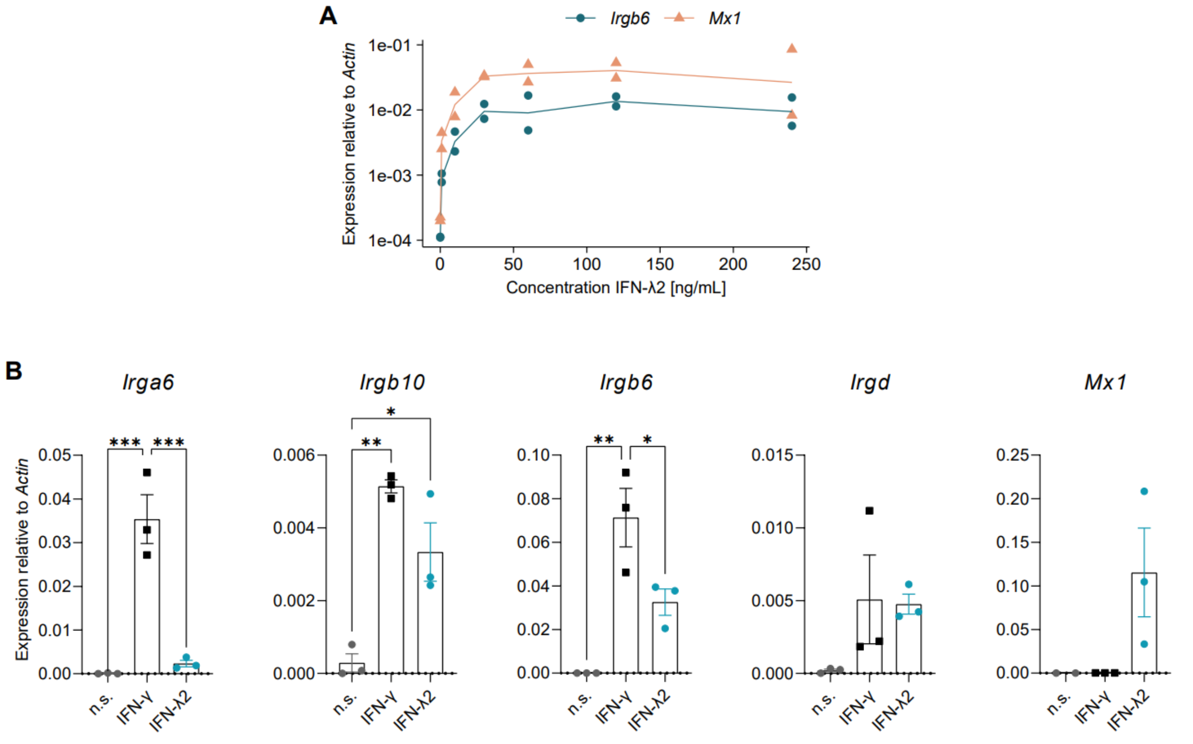


**Supplementary Figure 6. IRG effectors are induced by IFN-γ and IFN-λ2 in mouse intestine ODMs. A.** A saturated IFN-λ2-mediated response is reached with concentrations higher than 30 ng ml$^{-1}$. 7 day-differentiated ODMs were stimulated with 0 to 240 ng ml$^{-1}$of IFN-λ2 for 4 h. Expression of *Mx1* and *Irgb6* was determined by qPCR. Data represent two biological independent experiments. **B.** Gene expression of several effector *IRG*s and *Mx1* normalized to actin after treatment with 60 ng ml$^{-1}$IFN-γ or IFN-λ2 for 4 h was determined by qPCR. All analyzed *IRG*s except *Irgd* are induced by IFN-γ whereas only *Irgb6* and *Irgb10* are induced by IFN-λ. *Mx1* served as a positive control for IFN-λ2 treatment. Data represent the mean and SEM of three independent experiments with two replicates each, *p ≤ 0.05, ***p* < 0.01, ****p* < 0.001, *****p* < 0.0001 determined by ANOVA with Tukey's multiple-comparison test.

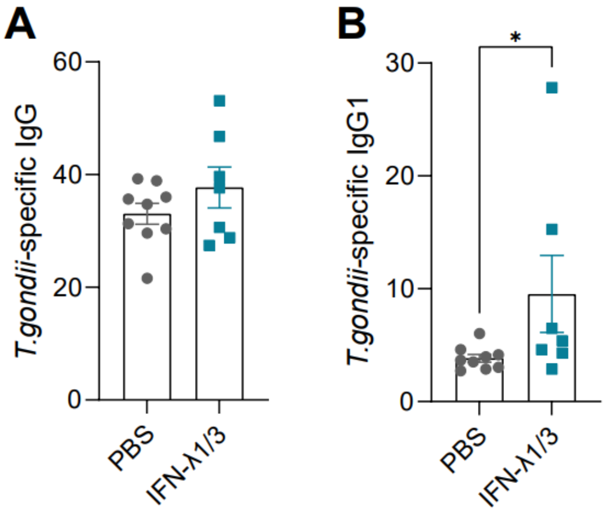


**Supplementary Figure 7. IFN-λ treatment of mice improves immune responses after oral *T. gondii* infection. A-F.** Mice were treated (i.p. injection) with 1 µg of IFN-λ1/3 or mock-treated with PBS/0.1 % BSA from day -1 to day 7 of oral *T. gondii* infection with 10 Pru-tdTomato-derived tissue cysts. *T. gondii*-specific IgGs and IgG1 were determined in serum by ELISA. Each dot represents an individual mouse, *p = 0.0390 determined by Mann Whitney test.

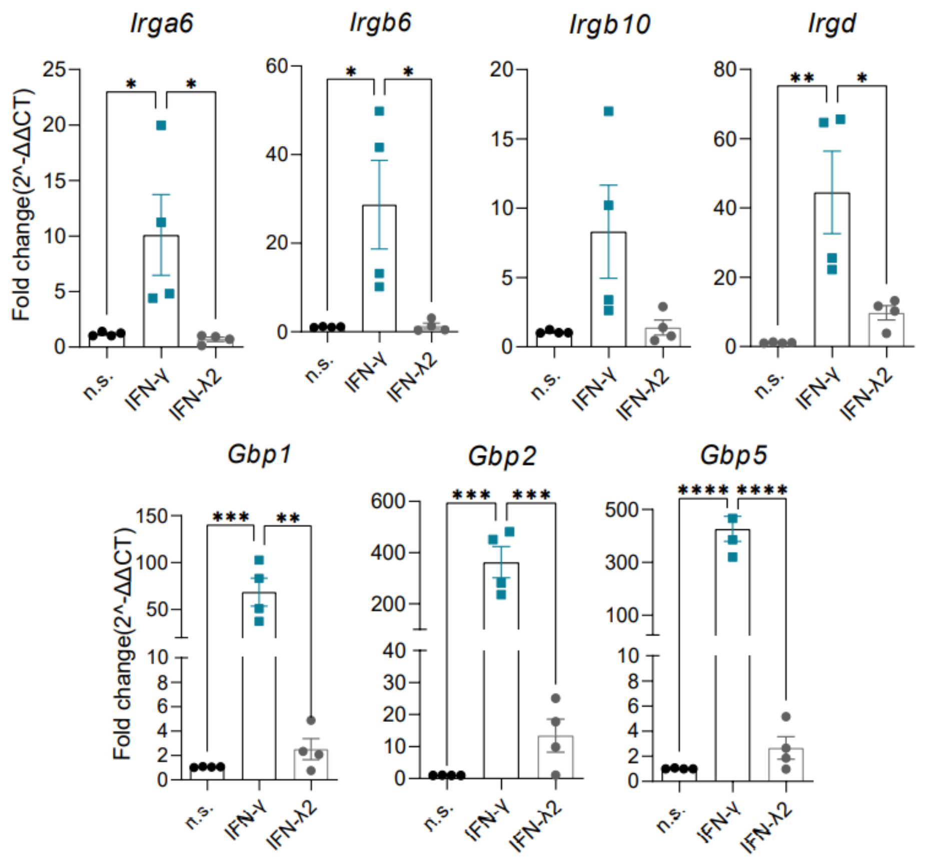


**Supplementary Figure 8. IRG and GBP proteins are significantly induced by IFN-γ but not by IFN-λ2 in neutrophils**. Bone marrow-derived neutrophils were primed for 4 h with 3 ng $ml^{-1}$ of IFN-γ or IFN-λ2 and the expression of *Irga6*, *Irgb6*, *irgb10*, *Irgd*, *Gbp1*, *Gbp2* and *Gbp5* was quantified by qPCR relative to *UbC*. Symbols represent means ±SEM of four independent experiments performed in triplicates, ****p < 0.0001, ***p < 0.001, **p < 0.01, *p < 0.05 determined by ANOVA with Tukey's multiple-comparison test.
